# The way less obvious: PIEZO1 supports apoptotic cell extrusion by optimizing tissue mechanical tension for homeostasis

**DOI:** 10.64898/2026.06.01.729446

**Authors:** Zoya Mann, Suzie Verma, Oscar Chen, Phoebe Dunbabin, Robert J. Ju, Changyuan Hu, Terry C.C. Lim Kam Sian, Edna C. Hardeman, Peter W. Gunning, Roger J. Daly, Kinga Duszyc, Kate Poole, Alpha S. Yap

## Abstract

Apical cell extrusion is a mechanical process that allows epithelia to eliminate apoptotic cells and prevent inflammation. Mechanosensitive ion channels are often invoked for their capacity to mediate rapid mechanical responses in dynamic morphogenetic processes. Here we report an unexpected strategy for PIEZO1 to support apoptotic extrusion. PIEZO1 inhibition blocks extrusion in cultured cells and zebrafish larvae. However, although PIEZO1 mediates calcium signals during the extrusion process, we show that extrusion is instead antagonized by increase in the preexisting mechanical tension of the epithelium when PIEZO1 is disrupted. Correcting enhanced pre-stress in PIEZO1-disrupted epithelia is sufficient to rescue apoptotic extrusion, even though it does not restore dynamic calcium signals. PIEZO1 supports mechanical homeostasis through a calcium/calcineurin-dependent pathway that protects MYPT1/myosin phosphatase from degradation to limit Myosin II activation. Therefore, PIEZO1 support the morphogenetic process of apoptotic extrusion through mechanical homeostasis.

## Introduction

Epithelia are mechanically active tissues that can exert tensile forces on their environment by coupling the contractile actin cytoskeleton to cell adhesions. These cell-generated forces participate in many morphogenetic processes during epithelial development and homeostasis ^1^. This is exemplified by the tissue mechanical changes that allow epithelia to eliminate apoptotic cells by apical extrusion (apoptotic extrusion; AE, for short ^2–4^). Epithelial cells become hypercontractile when they undergo apoptosis, which increases tension at adherens junctions (AJs) between the apoptotic cell and its immediate neighbours ^5^. This engages dynamic mechanical responses in those neighbouring cells. Notably, increased actomyosin contractility at the cortices immediately adjacent to the apoptotic cells allows neighbouring cells to exert force and expel the apoptotic cell from the epithelium ^3,6^. Thus, AE is driven by dynamic mechanical responses to the local force imbalance that apoptosis creates in the epithelium. Characteristically, these responses are local changes that occur within a few cell diameters of the apoptotic cell ^7,8^.

The dynamic mechanical events of AE do not take place in isolation. Instead, they occur within a mechanical landscape where the actomyosin cytoskeleton constantly exerts force on AJs ^9^ to generate a pre-existing tissue tension (tensile pre-stress). In morphologically quiescent epithelia these tensile forces are in balance, so that net cell movement does not occur. However, global tensile pre-stress can influence the efficacy of apical cell extrusion. In particular, increased tensile pre-stress in the epithelium antagonizes apical extrusion. This applied when extrusion was elicited by apoptosis ^10,11^ or expression of oncogenes ^12^; and when pre-existing tension was increased by diverse maneuvers, such as expression of a constitutively-active Myosin II regulatory light chain (MRLC) transgene ^10^ and depletion of caveolae ^12^. Thus, the efficacy of AE depends on pre-existing global tissue mechanics, as well as on the local dynamic events elicited by apoptosis.

Their critical role implies that tissue mechanics must be tightly regulated in epithelia for effective cell extrusion to occur. Of note, a growing number of mechanotransduction pathways are implicated in apical extrusion ^13,14^. Importantly, these regulatory pathways can impinge on different aspects of tissue mechanics. In one instance, a tension-sensing apparatus anchored to E-cadherin plays a key role in activating RhoA for the dynamic contractile changes that drive AE ^3^. This mechanism uses Myosin VI to detect increased tension applied to E-cadherin adhesions from apoptotic cells ^3,15^. In contrast, mechanosensitive caveolae support apical extrusion by regulating mechanical pre-stress in the epithelium ^12^. Thus, the distinct roles that dynamic and background global mechanics play in apical extrusion may reflect the impact of different mechano-regulatory pathways.

Mechanosensitive ion channels (MSCs) are attractive candidates to regulate epithelial mechanics ^16^, especially during dynamic morphogenetic events. The capacity for MSCs to respond rapidly to mechanical stimuli would place them well to coordinate rapid cellular responses for tissue rearrangements ^17–19^. Indeed, PIEZO1 and other MSCs have been implicated in apical cell extrusion ^20,21^, potentially via intracellular calcium signaling (Ca_i_) signaling ^22,23^, which is transiently activated within neighbour cells as the dynamic events of extrusion occur ^24^. However, the cellular pathways that might allow PIEZO1 to support extrusion are poorly understood. Nor is it known whether PIEZO1 functions as a primary force sensor of the dynamic mechanical changes that elicit extrusion, or whether it exhibits a basal activity that supports AE by acting more globally to control tissue pre-stress and mechanical homeostasis of the epithelium. In this study we combined genetic deletion of PIEZO1 with the use of PIEZO inhibitors to establish that PIEZO1 is necessary for AE *in vitro* and *in vivo*. We further show that PIEZO1 exerts this impact principally through the pre-existing global mechanical homeostasis of the epithelium. This function is achieved by a previously unrecognized pathway, where calcium/calcineurin protects Myosin phosphatase from degradation, thereby preventing excess activation of epithelial tissue tension.

## Results

We tested the effect of PIEZO1 on AE using MCF7 mammary epithelial cells and the periderm of zebrafish larvae. In both systems AJs assemble a contractile actomyosin cortex and display mechanical tension ^25–27^. qRT-PCR confirmed that PIEZO1 was expressed in MCF7^WT^ cells (Fig S1A), so we deleted it by CRISPR/Cas9 genome-editing (PIEZO1^KO^ or P1-KO). Deletion was confirmed by qRT-PCR and Western analysis (Fig S1B,C). PIEZO2 was not expressed in either wild-type (WT) cells or PIEZO1^KO^ cells (Fig S1A,B). Functional loss of PIEZO1 was tested by visualizing intracellular calcium (Ca_i_) signaling with Calbryte™520-AM. Baseline Ca_i_ levels were similar in both WT and PIEZO1^KO^ cells and were increased by the PIEZO activator Yoda1 ^28^ in WT but not PIEZO1^KO^ cells (Fig S1D,E). Conversely, GsMTx4, which inhibits PIEZO1 and other MSCs ^29–31^, decreased Ca_i_ levels in WT but not PIEZO1^KO^ cells (Fig S1D,E). As a further test of function, we tested for transient increases in Ca_i_ that have been reported to occur in a PIEZO1-dependent fashion in the epithelium around apoptotic cells ^21,23,24^. Consistent with this, we observed a brief (~60 sec), wave-like increase in Ca_i_ in the surrounding epithelium when apoptosis was induced by laser-irradiating individual cell nuclei (Fig S1F-H, Video S1). This effect was lost with PIEZO1^KO^ (Fig S1F-H, Video S1). Together, these results confirmed that KO cells were functionally deficient for PIEZO1, and this was without evident compensation.

### PIEZO1 is necessary for apoptotic extrusion

To test the role of PIEZO1 in AE, confluent MCF7 monolayers were treated with the topoisomerase inhibitor etoposide (500µM, 24h; Fig 1A). Western blot analysis for cleaved caspase-3 showed that apoptosis was induced to similar degrees in WT and in PIEZO1^KO^ lines (Fig S2A). Almost all apoptotic cells immunostaining for cleaved caspase-3 underwent AE in WT monolayers (Fig 1A). In contrast, AE was substantially reduced in the PIEZO1^KO^ monolayers (Fig 1A). Similarly, PIEZO1^KO^ significantly reduced the extrusion response when apoptosis was induced by nuclear laser irradiation (Fig 1B,C, Video S2). AE induced by etoposide was also inhibited when GsMTx4 was applied to MCF7^WT^ and Caco2^WT^ colon epithelial cells (Fig 1D, Fig S2B).

**Figure 1.**
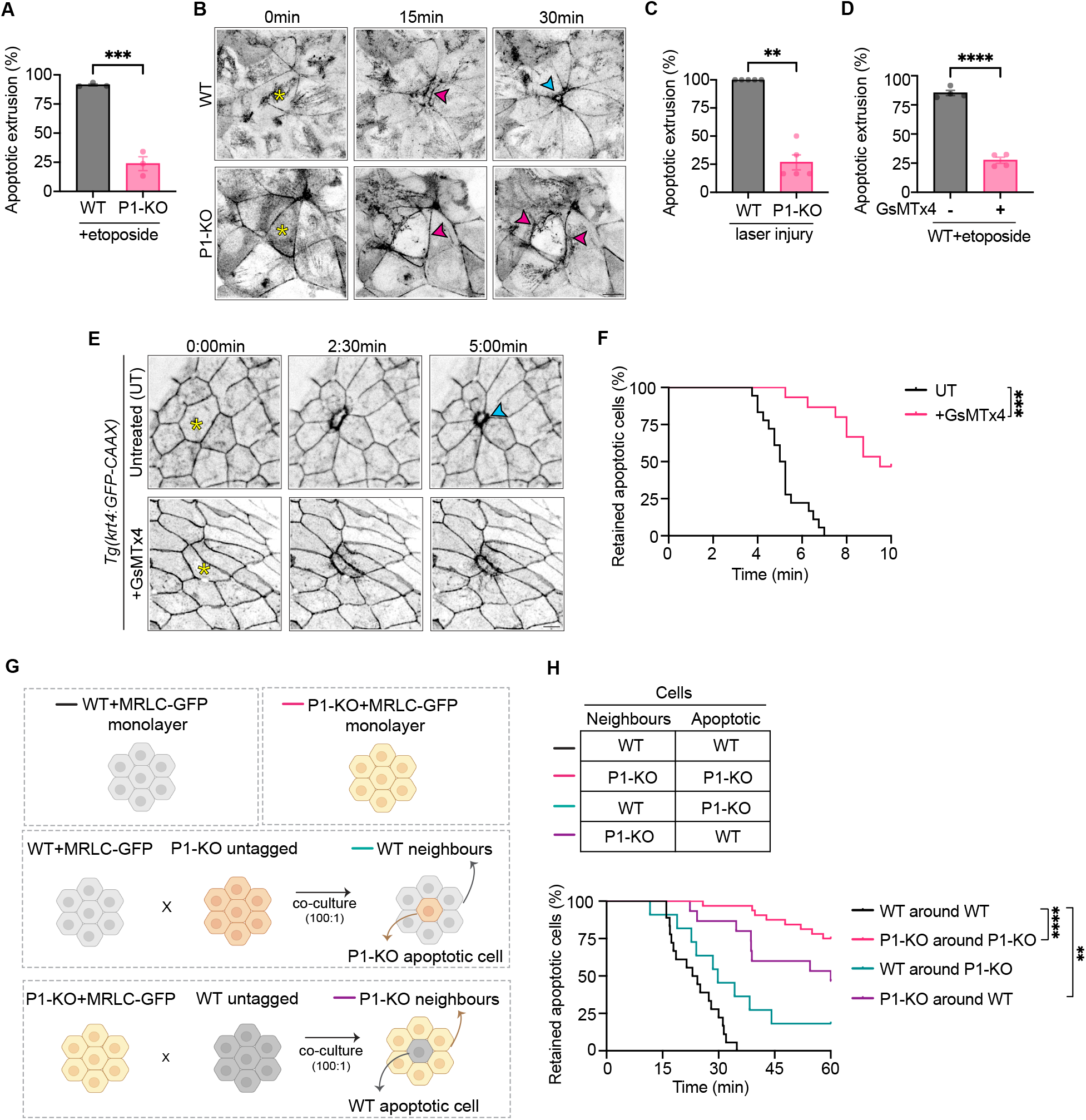
PIEZO1 is necessary for apoptotic cell extrusion. (A) Quantification of apoptotic extrusion (AE) efficiency in etoposide treated (500µM, 24h) wild-type (WT) and PIEZO1-knockout (P1-KO) monolayers. (B-C) Representative images (B) and quantification (C) of AE after laser-induced induction of apoptosis in WT and P1-KO monolayers. Yellow asterisk: cell selected for laser injury; magenta arrowheads: myosin ring formation at dead-neighbour cell interface, as visualised by MRLC-GFP; blue arrowhead: junctional closure underneath extruded apoptotic cell. (D) AE efficiency induced with etoposide (500µM, 24h) in WT monolayers with or without GsMTx4 pre-treatment (2.5µM). (E-F) Effect of GsMTx4 (5µM, 16h) on laser-induced AE in the periderm of *Tg(krt4:GFP-CAAX)* zebrafish larvae: representative frames (E) and quantification of time course (F). Yellow asterisk: cell selected for laser injury; blue arrowhead: junctional closure underneath extruded apoptotic cell. (G) Schematic outlining the strategy for growing mosaic monolayer cultures of WT and/or P1-KO cells. Untagged or MRLC-GFP expressing WT or P1-KO were either cultured as homogenous monolayers or co-cultured as mosaics (1:100 ratio) and grown to confluency before inducing laser-mediated cell injury. (H) Quantification of AE efficiency of laser-injury induced apoptosis in WT and P1-KO mosaic co-cultures. Untreated controls were treated with respective drug vehicles. Scale bars: 10µm. XY panels are maximum projection views of all z-stacks. All data are means ± SEM. **p<0.01, ***p<0.001, ****p<0.0001 calculated from n≥3 independent experiments analysed with unpaired Student’s *t* test (A,D), Mann-Whitney test (C), or two-way ANOVA (F,H).

We then induced apoptosis in zebrafish peridermal cells by nuclear laser irradiation, using a transgenic line that expresses a membrane marker to mark cell borders (*Tg(krt4:GFP-CAAX*); Fig 1E,F, Fig S2C-E, Video S4). Apoptotic cells underwent rapid apical extrusion in the periderm, but this effect was substantially compromised by the PIEZO inhibitors GsMTx4 (Fig 1E,F, Video S4) or Gd^3+ 20^ (Fig S2D,E, Video S4). Together, these results indicate that PIEZO1 is necessary for effective AE both *in vitro* and *in vivo*, across species.

### Apoptotic extrusion requires PIEZO1 in neighbour cells

Apoptotic extrusion is a cell non-autonomous process that requires active changes in both the apoptotic cell and in its proximate neighbours (“neighbour cells” for short) ^2,7^. We therefore performed mixing experiments with chimeric MCF7 monolayers to identify which of these populations required PIEZO1 for effective AE. Here, minorities of WT or PIEZO1^KO^ cells were surrounded by the alternate cell type and laser nuclear irradiation was used to induce apoptosis in the minority cells (Fig 1G). As predicted, AE was efficient when both apoptotic cells and their neighbours were WT and was substantially compromised when both subpopulations were PIEZO1^KO^ (Fig 1H, Video S2). Extrusion also proceeded effectively when the apoptotic cells were PIEZO1^KO^, but the neighbours were WT (Fig 1H, Fig S2F, Video S3). However, AE was substantially compromised when the neighbours were PIEZO1^KO^, even if the apoptotic cell was WT (Fig 1H, Fig S2F, Video S3). While subtle effects of PIEZO1 in the apoptotic cell cannot be excluded, these data indicated that effective AE requires PIEZO1 functionality in the neighbour cells.

To assess how PIEZO1^KO^ affected dynamic neighbour cell responses during AE, we visualized the cytoskeletal mechanisms that drive the extrusion process. Specifically, neighbour cells assemble a contractile cortex at the apoptotic:neighbour interface ^3,6,32^ which is necessary for effective extrusion in MCF7 cells ^3^. Therefore, we used live-cell imaging of myosin regulatory light chain (MRLC-GFP) to identify if the non-muscle myosin II (NMII) force-generator was compromised by PIEZO1^KO^ in neighbour cells. As previously observed, extrusion in WT monolayers was distinguished by the assembly of a Myosin II-rich contractile ring around apoptotic cells induced by laser irradiation (Fig 1B, Video S2). Surprisingly, prominent rings also assembled in PIEZO1^KO^ cells. Mixing experiments confirmed that KO neighbour cells effectively assembled contractile rings (Fig S2F, Video S3). This suggested that the extrusion defect observed with PIEZO1^KO^ was not due to contractile failure in neighbour cells.

### Tensile monolayer pre-stress is increased by PIEZO1^KO^

In considering alternative reasons for PIEZO1^KO^ to perturb AE, we were struck to observe that MRLC-GFP was enhanced at AJ throughout PIEZO1^KO^ monolayers, even away from the apoptotic cells (Fig 1B). This suggested that epithelial contractility might be elevated by PIEZO1^KO^, separate from the extrusion process itself.

Consistent with this notion, immunostaining revealed that activated Myosin II (pMRLC) was increased at AJs throughout PIEZO1^KO^ cell monolayers (Fig 2A-D), even without inducing apoptosis. Increased cellular levels of pMRLC were confirmed by Western analysis (Fig 2A,B); however, total MRLC levels were not elevated (Fig S3A), suggesting that activity of NMII was preferentially increased at AJs between PIEZO1^KO^ cells (Fig 2C,D). This was supported by GsMTx4 treatment (2.5µM, 15min), which increased activated Myosin II levels in MCF7^WT^ (Fig S3B,C,F,G) and Caco2^WT^ monolayers (Fig S3J-M). This was also evident *in vivo*, as GsMTx4 (5µM, 16h) increased total pMRLC in zebrafish larvae (Fig 2E,F) and this was apparent at AJs in the periderm (Fig 2G,H). Together, these findings indicate that PIEZO1 can antagonize contractility in epithelial sheets. This was confirmed using the PIEZO agonist, Yoda1 (25µM, 15min), which decreased levels of active NMII in MCF7^WT^ monolayers (Fig S3D,E).

**Figure 2.**
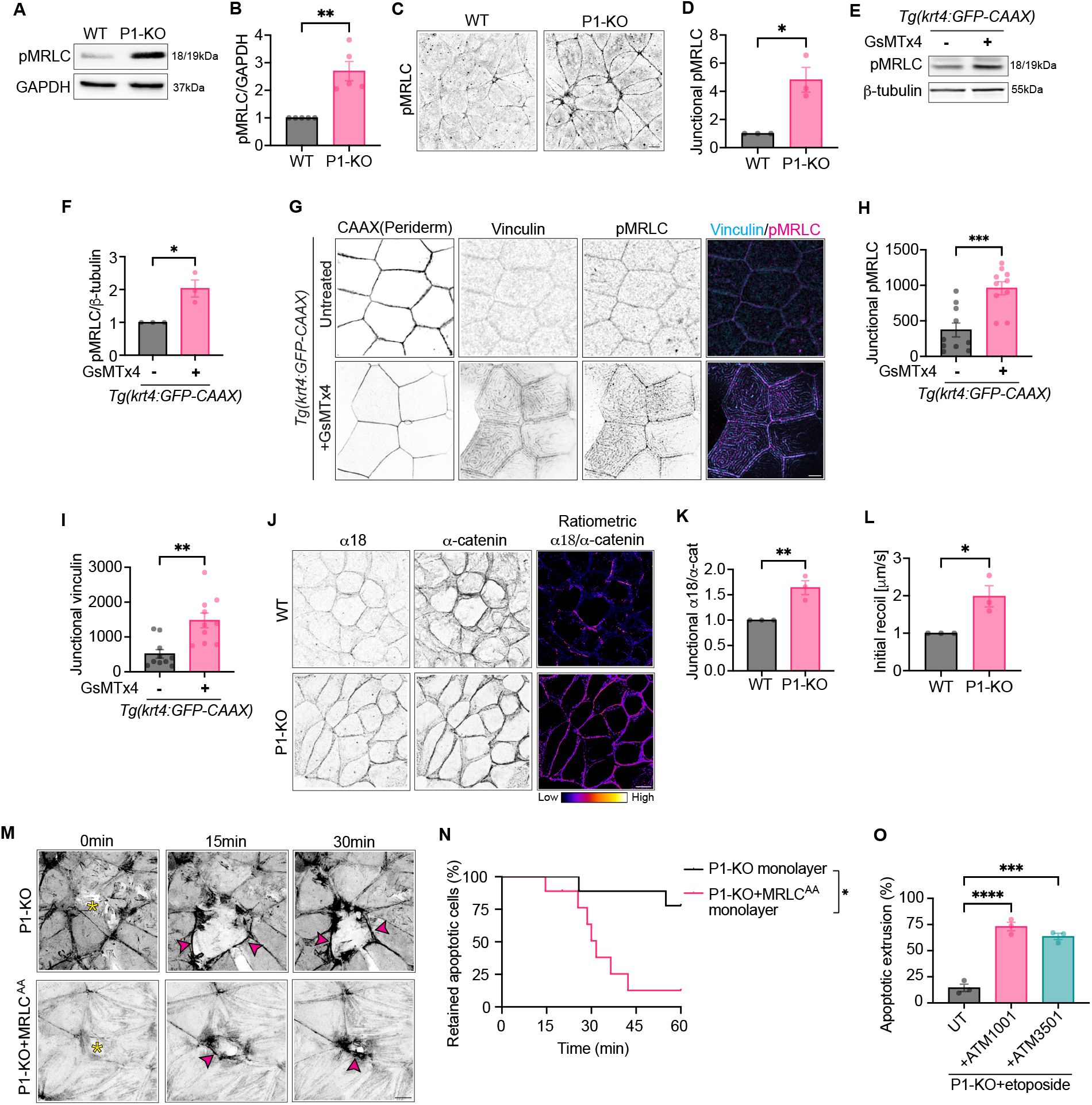
PIEZO1 disruption increases epithelial tissue tension. (A-B) Total phospho-myosin^S19^ (pMRLC) levels in WT and P1-KO cells: representative immunoblots (A) and quantification (B). (C-D) Representative immunofluorescent images (C) and quantification (D) of pMRLC localised at junctions of WT and P1-KO confluent monolayers. (E-F) Representative immunoblot (E) and quantification of total pMRLC levels (F) detected in *Tg(krt4:GFP-CAAX)* zebrafish larvae either untreated or pre-treated overnight with GsMTx4 (5µM, 16h). (G-I) Representative immunostained images (G) and quantification of pMRLC (H) and vinculin (I) localised to cellular junctions as visualised in periderm of *Tg(krt4:GFP-CAAX)* zebrafish larvae either untreated or pre-treated overnight with GsMTx4 (5µM, 16h). (J-K) Representative immunofluorescent images (J) of tension-sensitive α-catenin epitope (α18) and total α-catenin levels in WT and P1-KO monolayers. Ratiometric images (J) and quantification (K) represent junctional intensity of α18 normalised to total junctional α-catenin. (L) Tension measured by initial recoil of 2-photon laser-ablated junctions in WT and P1-KO monolayers. (M-N) Representative frames (M) and quantification (N) of live imaging of laser-induced apoptosis in P1-KO monolayers, compared to P1-KO monolayers expressing phosphodeficient myosin mutant, MRLC^AA^. Yellow asterisk-cell selected for laser injury; magenta arrowheads-myosin ring formation at dead-neighbour cell interface, as visualised by MRLC-GFP (in P1-KO) or MRLC^AA^-GFP (in P1-KO+MRLC^AA^). (O) Quantification of AE efficiency in etoposide treated (500µM, 24h) P1-KO monolayers in either untreated controls, or pre-treated with tropomyosin inhibitors (ATM1001, ATM3501; 2.5µM of each, 8h). Untreated controls were treated with respective drug vehicles. Scale bars: 10 µm. XY panels are maximum projection views of all z-stacks. All data are means ± SEM. *p<0.05, **p<0.01, ***p<0.001, ****p<0.0001 calculated from n≥3 independent experiments analysed with one-sample *t* test (B,D,F,K,L), unpaired Student’s *t* test (H,I), one-way ANOVA (O), or two-way ANOVA (N).

Activation of Myosin II would be expected to enhance mechanical tension in epithelia with deficient PIEZO1 function. To test this prediction, we first stained for a cryptic epitope of α-catenin (α18 mAb) that is revealed upon application of molecular tension to cadherin adhesion complexes ^33^. α18 mAb levels, corrected for total α-catenin, were increased at the AJs between PIEZO1^KO^ MCF7 cells (Fig 2J,K) and in both MCF7^WT^ and Caco2^WT^ monolayers treated with GsMTx4 (Fig S3H,I,N,O). This implied that mechanical tension was increased at AJs in PIEZO1-disrupted monolayers. As a more direct, biophysical measure of junctional tension we then measured the immediate velocity of recoil after AJs were cut with a laser ^34,35^. Indeed, PIEZO1^KO^ increased the initial recoil velocity of AJs, confirming increased junctional tension (Fig 2L).

To test if PIEZO1 also influences AJ tension *in vivo*, we stained for junctional vinculin in the zebrafish periderm. Vinculin is recruited to AJs in a tension-sensitive manner. We found that junctional vinculin was increased in GsMTx4-treated larvae (Fig 2G,I). Thus, apoptosis in PIEZO1-inhibited epithelia occurs against a background of increased tissue tension (tensile pre-stress).

### Elevated pre-stress antagonizes apoptotic extrusion when PIEZO1 is disrupted

Elevated tensile pre-stress antagonizes the ability of epithelia to mount an effective AE response ^10^. Therefore, we considered the possibility that increased pre-stress was responsible for compromising AE when PIEZO1 function is disrupted. We tested this hypothesis by reducing mechanical pre-stress in PIEZO1^KO^ cells before induction of apoptosis. For this we antagonized contractile tension throughout monolayers either by expression of the phospho-defective MRLC^AA^ mutant ^10^ or by treatment with the tropomyosin inhibitors (TPMis) ATM1001 and ATM3501 (2.5µM, 15min), which compromise the organization of NMII into contractile units^36^ (Fig S3P,Q). Both these maneuvers increased the proportion of apoptotic cells that underwent apical extrusion in MCF7 PIEZO1^KO^ monolayers (Fig 2M-O, Video S5). Thus, correction of global tissue mechanics could restore AE despite PIEZO1^KO^.

Together, these observations suggested that disrupting PIEZO1 inhibited AE by increasing the overall tensile pre-stress in the epithelial monolayer. However, Ca_i_ signaling in neighbours can also contribute to AE ^23,24^. As multiple mechanosensitive mechanisms support Ca_i_ signaling ^23,24^, it was possible that relieving the overall level of tension in PIEZO1^KO^ epithelia might restore Ca_i_ signaling by other mechanisms. However, the Ca_i_ response remained compromised in PIEZO1^KO^ cells treated with TPMi under conditions that improved AE efficiency (Fig S1F-H, Video S1). Therefore, restoration of the dynamic Ca_i_ signal was not responsible for restoring AE when pre-stress in the monolayer was ameliorated. This finding reinforces the notion that AE was disrupted by aberrant tensile pre-stress in the PIEZO1^KO^ monolayers.

### Myosin Phosphatase depletion increases tissue tension in PIEZO1^KO^

We then sought to understand how PIEZO1 regulated tensile pre-stress in epithelia. Myosin II activity is controlled by the balance between stimulatory pathways that phosphorylate the MRLC (Ser^18^, Thr^19^), countervailed by Myosin Light Chain Phosphatase (MLCP) that dephosphorylates these residues^37^. Formally, the enhanced MRLC phosphorylation evident in PIEZO1^KO^ cells could be due to an increase in NMII stimulatory pathways and/or a decrease in MLCP activity. RhoA signaling is a major stimulus for Myosin II at AJs ^25,38^. We predicted that RhoA activity would be elevated if changes in this pathway were responsible for the increased tissue tension of PIEZO1^KO^ monolayers. However, GTP-RhoA levels were reduced in PIEZO1^KO^ cells (Fig 3A,B), rather than being increased. Although we cannot exclude subtle effects of PIEZO1 on NMII stimulators, this result prompted us to focus on MLCP.

**Figure 3.**
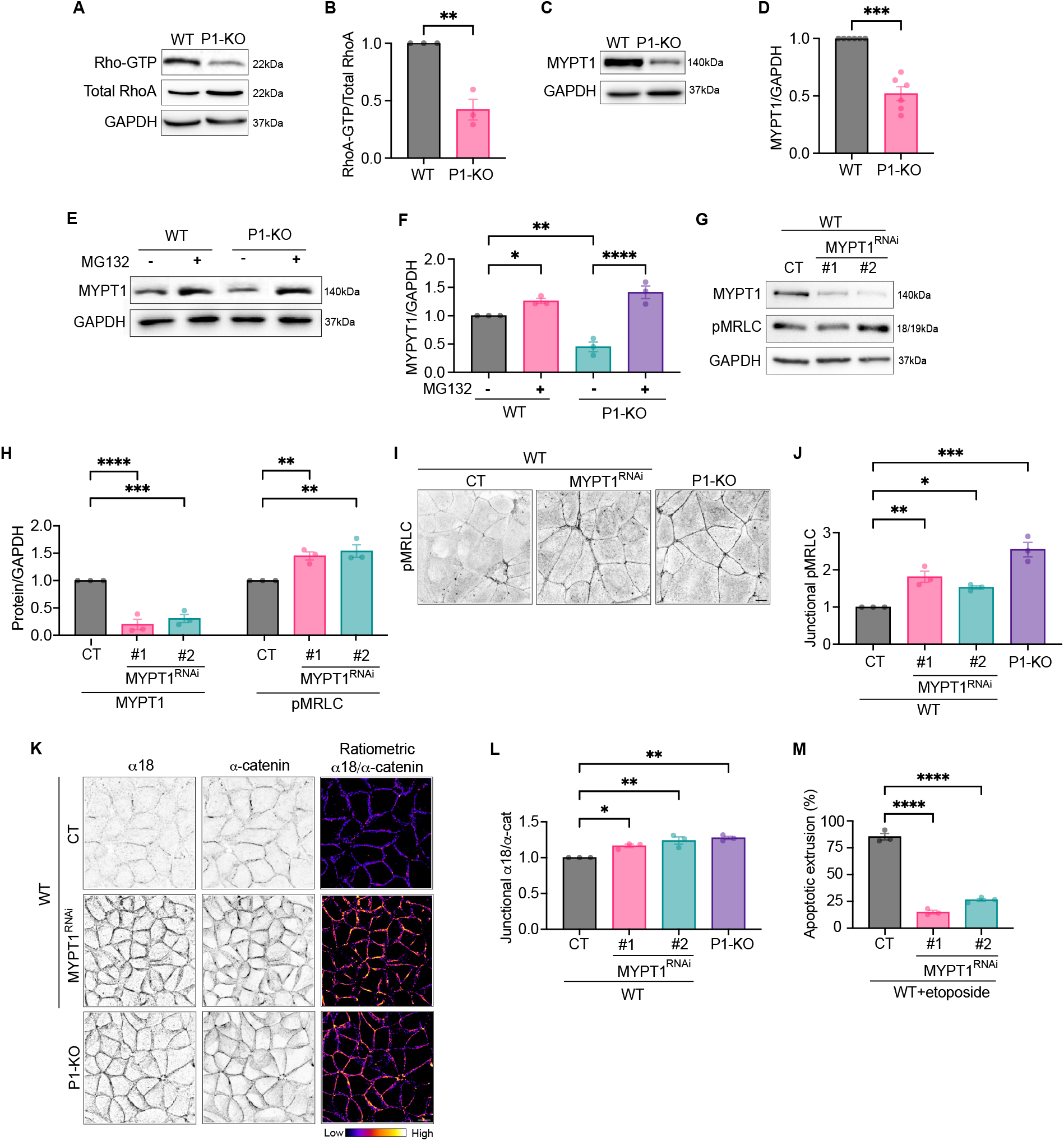
Myosin phosphatase is downregulated in PIEZO1^KO^ monolayers. (A-B) Active GTP-RhoA in WT and P1-KO cells: representative immunoblot (A) and quantification normalised to total RhoA protein levels (B). (C-D) Myosin phosphatase-regulatory subunit, MYPT1, levels in WT and P1-KO cells: representative immunoblot (C) and quantification (D). (E-F) Effect of proteosome inhibitor MG132 (50µM, 6h) on MYPT1 protein levels in WT and P1-KO cells: representative immunoblot (E) and quantification (F). (G-H) RNAi depletion of MYPT1 in WT cells and effect on pMRLC: representative immunoblots (G) and quantification (H) (48h post-MYPT1 knockdown). (I-J) Effect of MYPT1^RNAi^ in WT on junctional pMRLC compared with P1-KO: representative immunofluorescent images (I) and quantification (D). (K-L) Effect of MYPT1^RNAi^ in WT compared with P1-KO on junctional tension measured by α-catenin epitope (α18) relative to total α-catenin levels: representative ratiometric images (K) and quantification (L). (M) Quantification of AE efficiency in etoposide treated (500µM, 24h) control and MYPT1-depleted (MYPT1^RNAi^) WT monolayers. Untreated controls were treated with respective drugs vehicles. Scale bars: 10µm. XY panels are maximum projection views of all z-stacks. All data are means ± SEM. *p<0.05, **p<0.01, ***p<0.001, ****p<0.0001 calculated from n≥3 independent experiments analysed with one-sample *t* test (B,D), one-way ANOVA (H,M) or two-way ANOVA (F,J,L).

The MLCP holoenzyme consists of three elements: the regulatory subunit MYPT1, which is responsible for targeting and activating the conserved PP1c catalytic subunit, and a p20 subunit of unknown function ^39,40^. We focused on MYPT1 in order to identify changes that might regulate Myosin phosphatase in MCF7 cells. Strikingly, we found that cellular MYPT1 levels were significantly reduced in PIEZO1^KO^ MCF7 cells (Fig 3C,D). This decrease was attributable to protein degradation, as it was reversed by the proteosomal inhibitor MG132 (50µM, 6h; Fig 3E,F). Reduced MYPT1 levels would be predicted to increase MRLC phosphorylation. To test this principle, we depleted cellular MYPT1 by siRNA in WT cells (Fig 3G,H). Indeed, pMRLC phosphorylation was enhanced in MYPT1^RNAi^ cells (Fig 3G,H) and this was evident at AJs (Fig 3I,J). Tensile pre-stress was also increased in MYPT1^RNAi^, as reported by increased α18 mAb staining at AJs (Fig 3K,L) and AE was compromised (Fig 3M). This phenotypic similarity suggested that reduced MYPT1/MLCP might have been responsible for compromising AE when PIEZO1 was disrupted.

If so, we predicted that restoration of MYPT1 would reverse the aberrant contractile and extrusion responses seen in PIEZO1^KO^ monolayers. We tested this by exogenously expressing full-length MYPT1^HA^ in PIEZO1^KO^ cells (Fig 4). Lentiviral transduction restored overall cellular MYPT1 in PIEZO1^KO^ monolayers to levels similar to those of MCF7^WT^ (Fig 4A). This was accompanied by correction of NMII activity, measured by overall (Fig 4A) and junctional pMRLC levels (Fig 4B,C), and reversal of AJ tension assayed by α18 mAb immunolabelling (Fig 4D,E). Importantly, the efficacy of AE was increased when MYPT1^HA^ was expressed in PIEZO1^KO^ cells (Fig 4F). Together, these findings identified enhanced turnover of MYPT1/MLCP as responsible for the enhanced tissue pre-stress that inhibits AE when PIEZO1 is disrupted.

**Figure 4.**
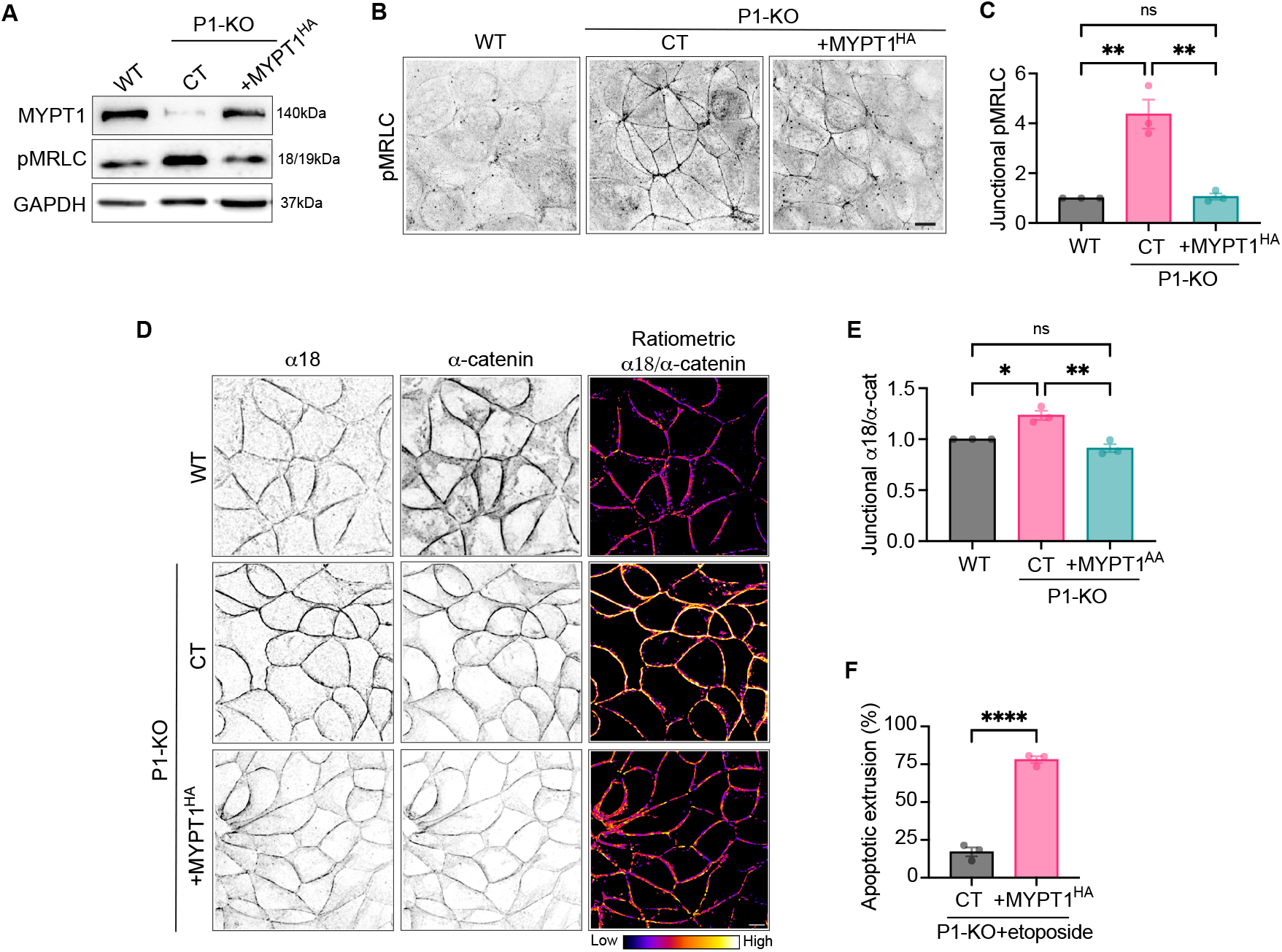
Exogenous myosin phosphatase expression reverts the mechanical phenotype and extrusion efficiency of PIEZO1^KO^ monolayers. (A) Exogenous expression of MYPT1^HA^ in PIEZO1^KO^: representative immunoblots of MYPT1 and pMRLC protein levels. (B-C) Effect of exogenous MYPT1^HA^ on junctional Myosin II activity (pMRLC immunostaining): representative immunofluorescent images (B) and quantification (C). (D-E) Effect on tension at AJ measured by tension-sensitive α-catenin epitope (α18) statining: representative images (D) and quantification (E) compared with total α-catenin. (F) Effect on AE efficiency with etoposide treatment (500µM, 24h). Scale bars: 10µm. XY panels are maximum projection views of all z-stacks. All data are means ± SEM. *p<0.05, **p<0.01, ****p<0.0001 calculated from n=3 independent experiments analysed with unpaired Student’s *t* test (F) or two-way ANOVA (C,E).

### MYPT1-subunit phosphorylation is increased by PIEZO1^KO^

To understand how MYPT1 turnover was increased when PIEZO1 was disrupted, we then probed for post-translational modifications that critically target proteins for proteosomal degradation. We immunoprecipitated MYPT1 from cells where protein levels were enhanced by blocking the proteosome with MG132 (50µM, 6h). Ubiquitination of MYPT1 was increased in PIEZO1^KO^ cells, consistent with turnover via the proteosome (Fig 5A,B).

**Figure 5.**
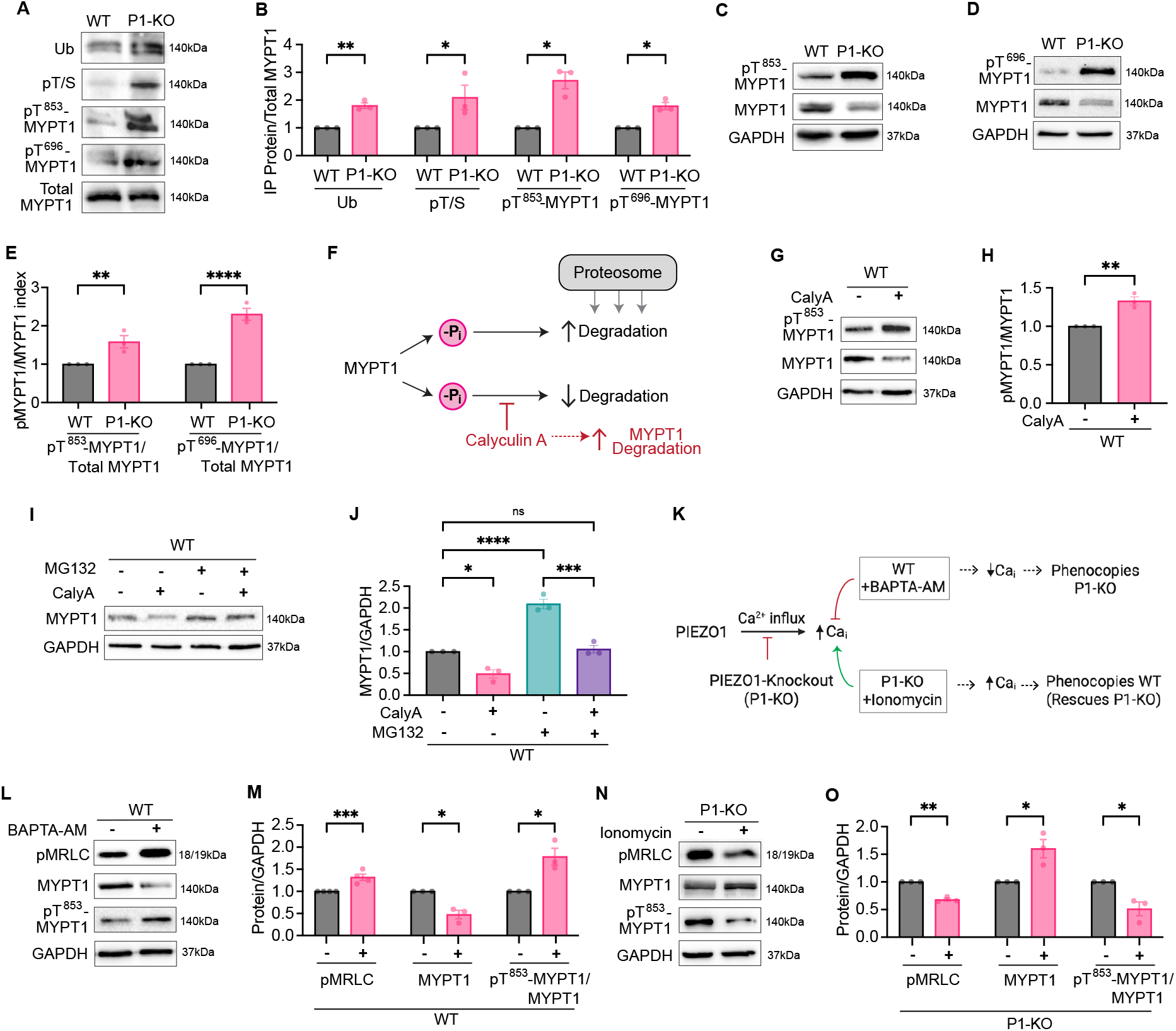
PIEZO1^KO^ affects calcium-sensitive post-translational modifications of MYPT1. (A-B) PIEZO1^KO^ induces post-translational modification (PTM) of MYPT1. MYPT1 immunoprecipitates probed for ubiquitin, phosphorylated threonine and serine (Thr/Ser or T/S), and specific pT^853^ and pT^696^ phospho-sites: representative immunoblots (A) and quantification of PTMs, normalized to total immunoprecipitated MYPT1 (B). (C-E) MYPT1 PTMs in whole-cell lysates of WT and P1-KO cells: representative immunoblots of pT^853^MYPT1 (C), pT^696^MYPT1 (D) and total MYPT1 (C,D), quantified relative to total MYPT1 (E). (F) Conceptual schematic for regulation of MYPT1 degradation by protein phosphorylation. The broad-spectrum Ser/Thr phosphatase inhibitor calcyulin A is predicted to increase MYPT1 phosphorylation and its degradation. (G-H) Effect of calyculin A (CalyA, 200nM, 15min) on pT^853^MYPT1 and total MYPT1 in MCF7^WT^ cells: representative immunoblots of whole-cell lysates (G) and quantification (H). (I-J) Effect of MG132 proteasome inhibitor (50µM, 6h) on cellular levels of MYPT1 in Ser/Thr-phosphatase-inhibited MCF7^WT^ cells (CalyA, 200nM, 15min): representative immunoblots of whole-cell lysates (I) and quantitation relative to GAPDH (J). (K) Conceptual design to test relationship between intracellular calcium and myosin regulation phenotype of PIEZO1^KO^. Intracellular calcium in WT cells was chelated with BAPTA-AM and calcium was increased in PIEZO^KO^ with ionomycin. (L-M) Effect of BAPTA-AM (10µM, 3h) on active myosin II (pMRLC), total MYPT1 and pT^853^-MYPT1 in MCF7^WT^ cells: representative immunoblots (L) and quantification (M). (N-O) Effect of ionomycin (2µM, 3h) on active myosin II (pMRLC), total MYPT1 and pT^853^-MYPT1 in P1-KO cells: representative immunoblots (N) and quantification (O). Untreated controls were treated with respective drugs vehicles. All data are means ± SEM. *p<0.05, **p<0.01, ***p<0.001, ****p<0.0001 calculated from n=3 independent experiments analysed with one-sample t-test (B,E,H,M,O) or one-way ANOVA (J).

Protein phosphorylation often collaborates with ubiquitination to target proteins for degradation ^41,42^. This possibility was suggested by finding that serine/threonine phosphorylation of MYPT1 was increased in PIEZO1^KO^ cells, compared with that from WT monolayers (Fig 5A,B). This was evident by probing MYPT1 immunoprecipitates with phospho-Ser/Thr antibodies and with antibodies that specifically recognize phosphorylation of Thr^696^ (pT^696^) and Thr^853^ (pT^853^), sites that have been implicated in inactivating MYPT1 upon phosphorylation (Fig 5A,B) ^43^. Enhanced pT^696^ and pT^853^ of endogenous MYPT1 was also evident in total PIEZO1^KO^ lysates, corrected for the lower MYPT1 protein levels in these cells (Fig 5C-E).

To test the principle that protein phosphorylation might promote MYPT1 degradation we treated MCF7^WT^ cells with the broad-spectrum phosphatase inhibitor, calyculin A ^44^(calyA; 200nM, 15min; Fig 5F). As expected, calyculin A increased MYPT1 phosphosphorylation (Fig 5G,H). This was accompanied by reduced levels of MYPT1, a change that was counteracted by MG132 (Fig 5F,I,J; Suppl Fig S4A-D). These data reinforced the notion that loss of protein phosphatase activity might target MYPT1 for degradation. Interestingly, calyculin A by itself did not further decrease MYPT1 levels in PIEZO1^KO^ cells (Fig S4C,D). Calyculin A enhanced NMII activity in MCF7^WT^ cells; however, it did not further increase the already elevated levels of pMRLC found in PIEZO1^KO^ cells (Fig S4A,B). Since calyculin A stimulates NMII by inhibiting myosin phosphatase ^44^, this finding suggests that myosin phosphatase activity is already substantially reduced by PIEZO1^KO^.

### PIEZO1-dependent calcium signaling regulates MYPT1 turnover

PIEZO1 mediates rapid, transient and localized flux of cations across the plasma membrane, including Ca^2+ 45^ and this localized activity can be generated by cellular contractility ^46,47^. Although loss of the acute transient calcium wave did not explain how PIEZO1 depletion inhibited AE, it was possible that basal PIEZO1 activity could influence the tensile pre-stress of the epithelium. Accordingly, we then manipulated Ca_i_ to test the principle of this hypothesis and its potential relationship with changes in MYPT1.

We first antagonized cellular calcium signaling in MCF7^WT^ cells with BAPTA-AM (10µM, 3h; Fig 5K-M), a cell-permeant pro-drug that chelates intracellular calcium when it is processed to release BAPTA in the cytoplasmic compartment (Fig S1D,E) ^48^. We predicted that if PIEZO1 were regulating epithelial contractility via calcium, then antagonizing intracellular calcium signaling should mimic the PIEZO1^KO^ phenotype even in MCF7^WT^ cells (Fig 5K). Indeed, we found that BAPTA-AM decreased MYPT1 levels and increased pMRLC in WT cells, consistent with the notion that downregulating MLCP enhances contractility (Fig 5L,M). This was accompanied by an increased in MYPT1 phosphorylation (pT^853^), corrected for the reduced level of total MYPT1 (Fig 5L,M). Thus, blocking calcium signaling altered myosin regulation in WT cells exactly as seen with PIEZO1^KO^.

Then, we stimulated calcium signaling in PIEZO1^KO^ cells using the calcium ionophore, ionomycin (2µM, 3h; Fig S1D,E). This tested the prediction that, if PIEZO1 were regulating epithelial contractility via calcium, then the KO phenotype would be reversed by stimulating calcium entry into the cell (Fig 5K). In support, ionomycin increased MYPT1 levels in PIEZO1^KO^ cells (Fig 5N,O), reinforcing the notion that its turnover was enhanced through loss of a calcium-signaling mechanism in PIEZO1-defective cells. This was accompanied by reduced phosphorylation (pT^853^) of MYPT1 (Fig 5N,O). Finally, ionomycin reduced activated NMII (pMRLC) from the elevated levels seen with PIEZO1^KO^ (Fig 5N,O).

Overall, these results support the concept that contractility is increased in PIEZO1-deficient cells because defective calcium signaling promotes MYPT1/MLCP turnover. In other words, PIEZO1-dependent calcium signaling protects MYPT1/MLCP from degradation. They further reinforced the hypothesis that a key targeting step might involve the phosphorylation status of MYPT1. Specifically, PIEZO1 would antagonize MYPT1 phosphorylation and degradation to preserve mechanical homeostasis.

### Calcineurin mediates mechanical homeostasis for AE

If PIEZO1-dependent calcium signaling were to antagonize phosphorylation of MYPT1, this implied that a calcium-dependent protein phosphatase might play a key role in protecting MYPT1 from degradation, to ultimately limit build-up of tissue tension. Interestingly, the calcium-dependent protein phosphatase, calcineurin, has been identified in controlling MYPT1 ^49^, providing a candidate for PIEZO1 to regulate epithelial contractility.

We tested this possibility using the calcineurin inhibitor Cyclosporine A (CsA; 50µM, 3h), predicting that if calcineurin mediates PIEZO1 signaling, then its inhibition would mimic the PIEZO1^KO^ phenotype (Fig 6A). Indeed, CsA increased MYPT1 phosphorylation (pT^696^) in MCF7^WT^ cells to levels similar to those seen with PIEZO1^KO^ (Fig 6B,E). This change in phosphorylation was accompanied by reduced levels of total MYPT1 protein (Fig 6C,E). Interestingly, CsA had little to no effect on the already aberrant levels of pMYPT1 and MYPT1 seen in PIEZO1^KO^ cells. This identified calcineurin as a dominant MYPT1 protein phosphatase that regulated its protein turnover.

**Figure 6.**
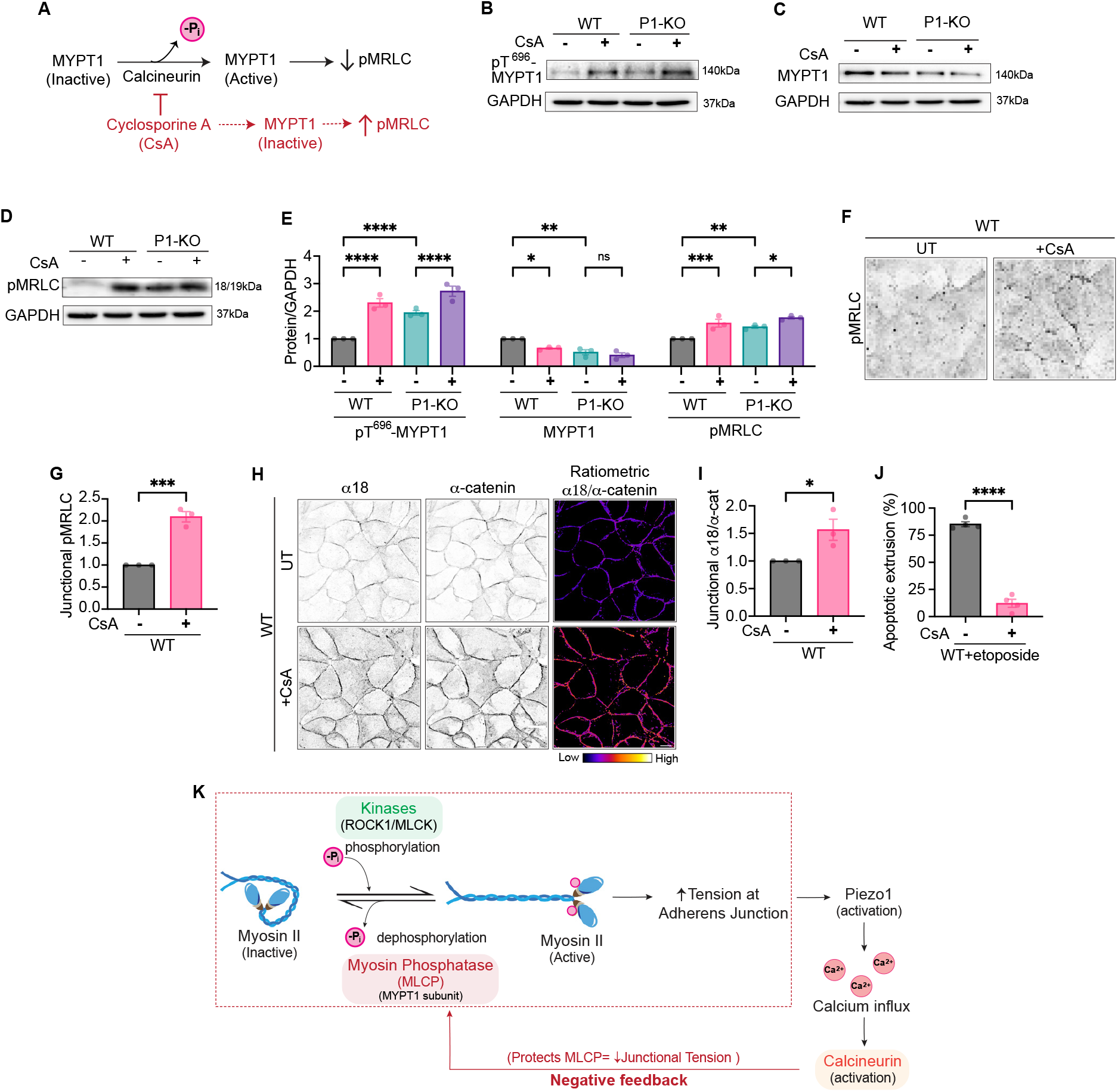
PIEZO1^KO^ affects calcium-sensitive post-translational modifications of MYPT1. (A-B) PIEZO1^KO^ induces post-translational modification (PTM) of MYPT1. MYPT1 immunoprecipitates probed for ubiquitin, phosphorylated threonine and serine (Thr/Ser or T/S), and specific pT^853^ and pT^696^ phospho-sites: representative immunoblots (A) and quantification of PTMs, normalized to total immunoprecipitated MYPT1 (B). (C-E) MYPT1 PTMs in whole-cell lysates of WT and P1-KO cells: representative immunoblots of pT^853^MYPT1 (C), pT^696^MYPT1 (D) and total MYPT1 (C,D), quantified relative to total MYPT1 (E). (F) Conceptual schematic for regulation of MYPT1 degradation by protein phosphorylation. The broad-spectrum Ser/Thr phosphatase inhibitor calcyulin A is predicted to increase MYPT1 phosphorylation and its degradation. (G-H) Effect of calyculin A (CalyA, 200nM, 15min) on pT^853^MYPT1 and total MYPT1 in MCF7^WT^ cells: representative immunoblots of whole-cell lysates (G) and quantification (H). (I-J) Effect of MG132 proteasome inhibitor (50µM, 6h) on cellular levels of MYPT1 in Ser/Thr-phosphatase-inhibited MCF7^WT^ cells (CalyA, 200nM, 15min): representative immunoblots of whole-cell lysates (I) and quantitation relative to GAPDH (J). (K) Conceptual design to test relationship between intracellular calcium and myosin regulation phenotype of PIEZO1^KO^. Intracellular calcium in WT cells was chelated with BAPTA-AM and calcium was increased in PIEZO^KO^ with ionomycin. (L-M) Effect of BAPTA-AM (10µM, 3h) on active myosin II (pMRLC), total MYPT1 and pT^853^-MYPT1 in MCF7^WT^ cells: representative immunoblots (L) and quantification (M). (N-O) Effect of ionomycin (2µM, 3h) on active myosin II (pMRLC), total MYPT1 and pT^853^-MYPT1 in P1-KO cells: representative immunoblots (N) and quantification (O). Untreated controls were treated with respective drugs vehicles. All data are means ± SEM. *p<0.05, **p<0.01, ***p<0.001, ****p<0.0001 calculated from n=3 independent experiments analysed with one-sample t-test (B,E,H,M,O) or one-way ANOVA (J).

Consistent with its inhibitory effect on MYPT1, CsA increased cellular and junctional NMII activity (Fig 6D-G). This effect was accompanied by increased mechanical tension at AJs reported with mAb α18 staining (Fig 6H,I). Importantly, blocking calcineurin inhibited AE in MCF7^WT^ cells (Fig 6J), as seen with PIEZO1^KO^. Together, these data identify calcineurin as a potential candidate to mediate PIEZO1/calcium signaling for mechanical homeostasis and AE in epithelia.

## Discussion

PIEZO channels have been implicated in many biological processes ^18,19,50^. In epithelial tissues, the impact of these MSCs ranges from stem cell turnover ^51^ to morphogenesis ^52,53^, and they mediate responses of cancer cells to mechanical stimuli ^54,55^. While the molecular and biophysical properties of PIEZO channels have been studied in detail, the challenge remains to understand how these molecular mechanisms elicit their functional consequences at the cell and tissue scales ^45^. Taken with recent reports ^21,23^, our data establish that PIEZO1 is necessary for effective AE. Unexpectedly, our experiments reveal that this functional impact reflects the operation of a countervailing pathway for global mechanical homeostasis, which promotes turnover of NMII activity to limit the aberrant build-up of epithelial tissue tension.

Apoptotic extrusion is a cell non-autonomous process, that requires dynamic mechanical responses of neighbour cells ^2,7^. Consistent with this, we found that PIEZO1 was predominantly required in neighbour cells to support AE. It was therefore attractive to postulate that PIEZO1 might participate in the dynamic cytoskeletal responses that allow neighbour cells to drive AE. This notion was supported by the transient, PIEZO1-dependent calcium waves that we and others ^21^ observed in neighbour cells shortly after induction of apoptosis. In one plausible model, PIEZO1 would have operated as a mechanotransducer that detected the local change in force induced by apoptosis to elicit contractile responses via signals such as calcium.

Instead, our data indicate that PIEZO1 supports AE by controlling the pre-existing mechanics of the epithelium. PIEZO1 inactivation by genetic deletion or inhibitor drugs increased the global tension of epithelia, both in cultured cells and in the zebrafish periderm, while the PIEZO activator Yoda1 inhibited NMII activity in WT cells. Increased tensile pre-stress antagonizes apical cell extrusion ^10,12^. Indeed, reversal of the tensile pre-stress restored the efficacy of AE in PIEZO1^KO^ cells. This seen with diverse maneuvers that reversed pre-stress, including directly relaxing the contractile apparatus with MRLC^AA^ and TPMi, and manipulating elements of the signaling pathway that allows PIEZO1 to modulate pre-stress. Importantly, AE was restored to PIEZO1^KO^ cells when tensile pre-stress was reversed by TPMi, even though the calcium waves were not restored. It is possible that PIEZO1-dependent Ca_i_ transients participate in some other aspect of the AE response. Nonetheless, our findings indicate that limiting pre-existing tissue tension is a dominant pathway for PIEZO1 to support AE. Therefore, PIEZO1 supported the dynamic process of AE by conditioning global tissue mechanics, rather than necessarily acting as a primary mechanosensor that is activated by the dynamic process itself.

How does PIEZO1 limit the build-up of tensile pre-stress in epithelia? Despite its being the best characterized regulator of epithelial tension ^56^, changes in RhoA did not explain the impact of PIEZO1 inhibition on epithelial contractility. Instead, we found that PIEZO1 supported epithelial mechanical homeostasis via myosin phosphatase, which inactivates NMII to balance the action of pro-contractile signals ^37,39^. MYPT1 levels were reduced when PIEZO1 was disrupted, reflecting increased proteosomal turnover, and MYPT1^RNAi^ alone mimicked the effect of targetting PIEZO1, making its loss a plausible explanation for the observed effects of PIEZO1 disruption. Importantly, expression of exogenous MYPT1 reverted tissue tension and restored AE efficiency to PIEZO1^KO^ cells. This causal linkage argues that PIEZO1 supports AE by limiting the turnover and degradation of MYPT1, thereby ensuring that Myosin II is dephosphorylated to prevent the build-up of contractility.

While PIEZO1 is a non-selective cation channel ^45^, calcium influx is a key consequence of PIEZO1 activation. Several lines of our data implicate a calcium-dependent signaling pathway that engages calcineurin to limit MYPT1 phosphorylation. First, direct manipulation of Ca_i_ closely paralleled the effects on MYPT1 and NMII activity seen with PIEZO1 disruption. Chelating intracellular calcium in WT cells mimicked PIEZO1^KO^ and, conversely, a calcium ionophore reverted contractile activation in PIEZO1^KO^ cells. Although PIEZO1 did not alter overall baseline Ca_i_ levels in our experiments, its activation at cell-substrate adhesions induced membrane-local flashes of Ca^2+ 46^ that may have been below the resolution of our assays.

Second, MYPT1 phosphorylation correlated inversely with its protein levels, a trend that was consistent across diverse maneuvers used in our experiments. Third, inhibiting calcineurin in WT cells exactly phenocopied the effect of PIEZO1 disruption, causing the phosphorylation and degradation of MYPT1, associated with increased epithelial tension and defective extrusion. Whether calcineurin directly or indirectly dephosphorylates MYPT1 remains to be determined. Phosphorylation of MYPT1 is known to promote cell contractility by inhibiting MLCP activity ^57^. Based on our findings, enhanced turnover of MYPT1 could synergize with enzymatic inhibition of MLCP to further increase contractility.

Together, these findings support a model where PIEZO1 participates in epithelial mechanical homeostasis via a Ca^2+^/calcineurin pathway that protects MYPT1/MLCP from being targeted for degradation and turnover (Fig 6K). By countervailing NMII activation, MYPT1/MLCP would limit the build-up of tissue tension, thereby facilitating AE. This implies that the MYPT1/MLCP limb is a regulated locus for mechanical homeostasis, not simply a default pathway. But why would a PIEZO1-dependent pathway be operating to control tension in epithelia that appear morphologically quiescent, where forces would be predicted to be balanced? Of note, even in mature epithelial monolayers contractility at AJs leads to transient cortical condensation ^9^, with the potential to generate small, unbalanced forces that could activate PIEZO1 at cell-cell contacts ^58–60^, as has been found for cell-substrate adhesions ^61 62 46^. Local activation of PIEZO1-Ca^2+^/calcineurin-MYPT1 signaling at cell-cell contaacts would then provide a negative feedback pathway that limits increase in junctional tension and thereby help ensure that forces are balanced in mature epithelia.

In conclusion, we have found that in epithelia PIEZO1 can participate in tissue mechanical homeostasis through a pathway that countervails the activation of cell contractility. This reinforces the concept that PIEZO1 can influence dynamic morphogenetic events by conditioning general tissue mechanics ^16^ as well as participating in those dynamic events. Indeed, our findings imply that for AE the impact on general tissue mechanics can be more critical than its acute effects. Of note, epithelial tension was increased by relatively short exposure to GsMTx4/Gd^3+^, but this was sufficient to inhibit AE. Therefore, morphogenetically-signficant changes in tissue mechanics can arise soon after changes in PIEZO1 activity. In future research, it will be important to test if the principles that we discovered for apoptosis apply to other forms of apical cell extrusion, and to other forms of epithelial morphogenesis.

## Limitations of the current study

We would highlight two areas for future study. First, more detailed biochemistry will be necessary to elucidate the molecular mechanisms of the pathway that we have described at the cell-biological level. These would include mapping post-translational modifications that may target MYPT1 for degradation. Second, PIEZO has also been reported to stimulate contractility ^63^, an effect that may be directly mediated by calcium signaling or indirectly via RhoA. However, in this case, PIEZO1 was more highly expressed in fibre cells, rather than epithelia, indicating that cell background may be a key determinant of the impact of PIEZO1 activation on contractility. How cells balance these effects against the tension-limiting pathway that we have identified remains to be determined. One possibility is that PIEZO1-activated calcium signaling can activate both stimulatory and inactivation limbs of the signaling network that controls NMII activity, via MLCK and calcineurin/MYPT1, respectively, depending on the localisation and magnitude of PIEZO1 activation, the specific cellular background or the specific mechanical context of the tissue. This flexibility could ensure turnover of NMII activity, to tune the contractile response in response to stimulatory inputs. Here it will be important to consider if PIEZO1 has different effects on contractility depending on the presence or absence of AJs. Cell-generated forces induce PIEZO1-mediated Ca^2+^ flashes that are transient and localized ^46^. So, PIEZO1-mediated signals may affect different balances of pro- and anti-contractile signaling, depending on whether they are activated at AJs or different parts of the cell.

## Acknowledgements

ZM, SV, RJ, KD and ASY were funded by the Australian Research Council (DP220103951, DP260103470) and the National Health and Medical Research Council of Australia (GNTs 2010704 and 2036946). RJJ is a recipient of a Cure Cancer ECR grant (CCAF2025-JU) and is an NHMRC Investigator (2042103). ASY is an Australian Laureate Fellow of the ARC (FL230100100).

PWG and ECH were funded by the Australian National Health and Medical Research Council (NHMRC) (APP2011770).

KP was funded by NHMRC (GNT1185021).

Imaging was performed in part at the IMB/ACRF Centre for Cancer Imaging, established with the support of the Australian Cancer Research Fund.

## Conflicts of Interest

Peter W. Gunning and Edna C. Hardeman own shares and are Directors of the board of the company, TroBio Therapeutics, that is developing drugs that target the functions of tropomyosins.

Alpha S. Yap is a member of the Editorial Board of Developmental Cell.

