## Supplementary Figures for "The way less obvious: PIEZO1 supports apoptotic cell extrusion by optimizing tissue mechanical tension for homeostasis"

**Figure S1.**

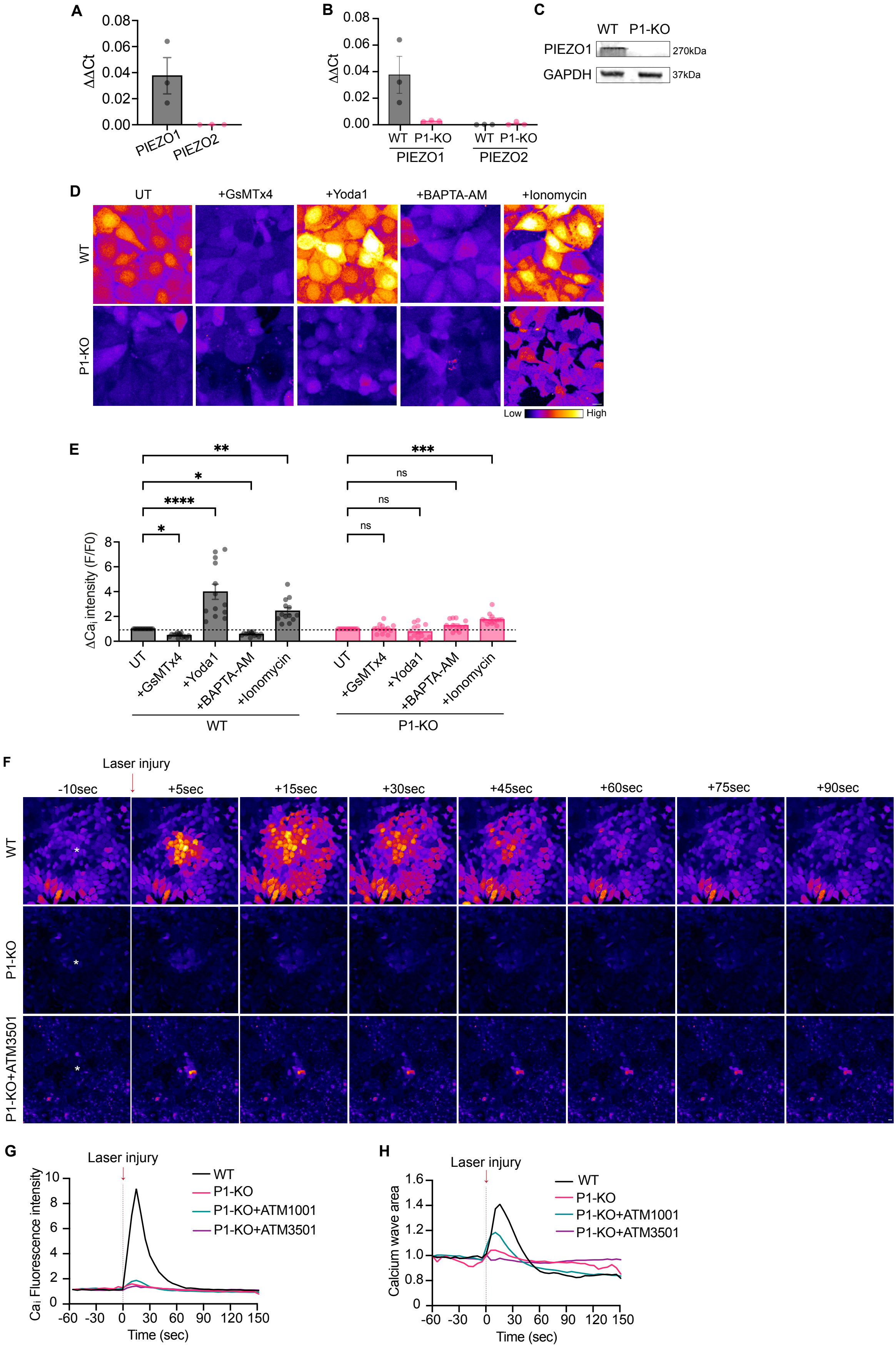

**Figure S1.**

(A) $\Delta\Delta C_t$ values of PIEZO1 and PIEZO2 mRNA expression detected via qRT-PCR as observed in $MCF7^{WT}$ cells.
(B) $\Delta\Delta C_t$ values of PIEZO1 and PIEZO2 mRNA expression detected via qRT-PCR in WT and P1-KO cells.
(C) Representative immunoblot showing PIEZO1 protein levels in WT and P1-KO cells.
(D-E) Representative images (D) and quantification (E) revealing foldchange in intracellular calcium ($\Delta Ca_i$) levels in GsMTx4 (2.5μM, 5min), Yoda1 (25μM, 5min), BAPTA-AM (10μM, 3h), or Ionomycin (2μM, 3h) treated WT and P1-KO cells, as compared to respective untreated controls.
(F-H) Selected stills (F) and quantification (G,H) from live-imaging of untreated WT and P1-KO, and P1-KO treated with tropomyosin inhibitors (ATM1001, ATM3501) (2.5μM of each, 8h), recording dynamic $Ca_i$ changes upon laser-injury induced apoptotic extrusion. Quantification reveals fold-change in $Ca_i$ fluorescence intensity across time (min) (G) and spread of calcium wave area post-laser injury (H). White asterisk- cell selected for laser injury.
Untreated controls were treated with respective drug vehicles. Scale bars: 10μm. XY panels are maximum projection views of all z-stacks. All data are means ± SEM. $p<0.05$, $^{**}p<0.01$, $^{***}p<0.001$, $^{****}p<0.0001$ calculated from $n\geq3$ independent experiments analysed with one-way ANOVA (E).

**Figure S2.**

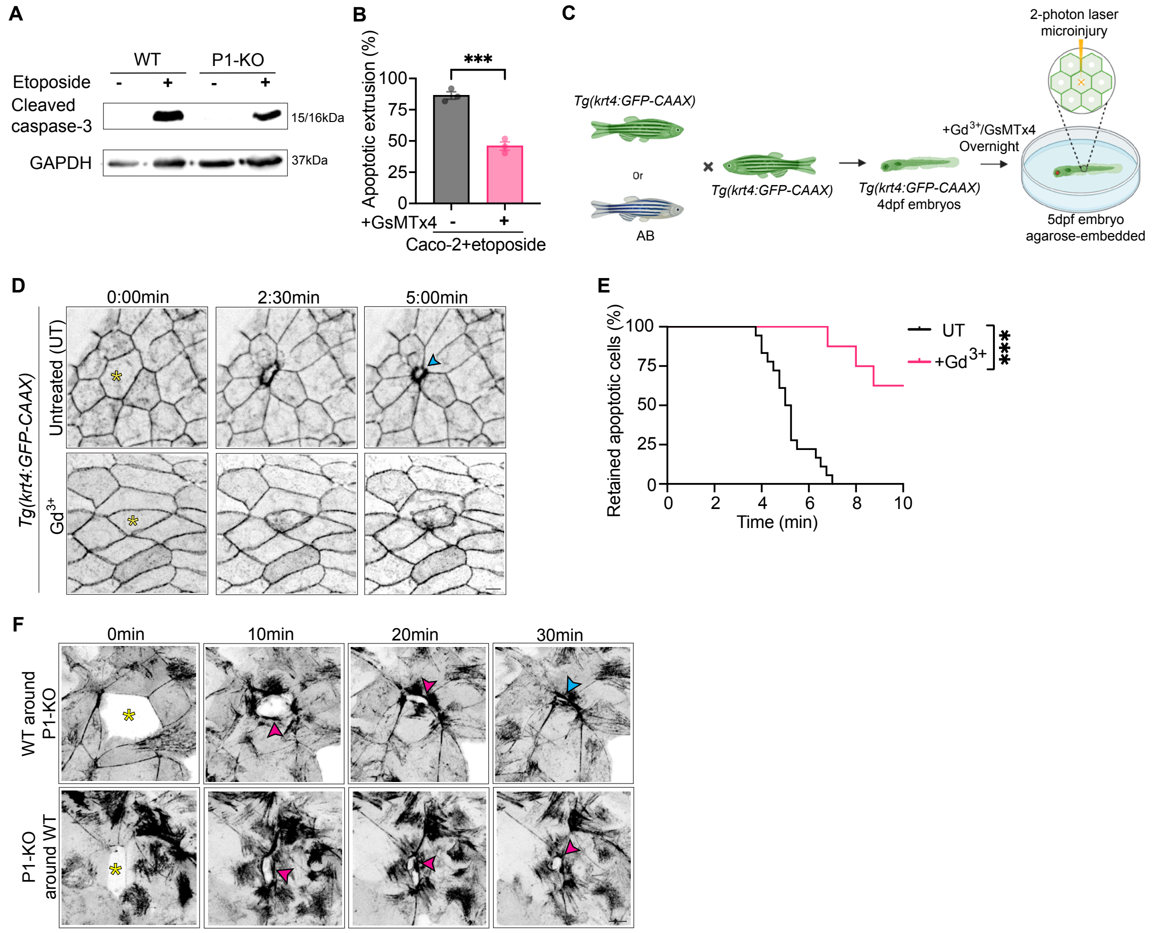


**Figure S2.**

(A) Representative immunoblot comparing percentage of apoptotic cells via cleaved-caspase-3 expression in untreated and etoposide-treated (500µM, 24h) WT and P1-KO monolayers.

(B) Quantification of apoptotic extrusion efficiency in etoposide treated (500µM, 24h) Caco-2 untreated and GsMTx4 pre-treated (2.5µM) monolayers.

(C) Schematic representing workflow of laser-induced cellular injury experiments done on zebrafish larvae. Briefly, 4dpf larvae were treated with GsMTx4 (5µM, 16h) or $Gd^{3+}$ (2.5µM, 16h) and embedded in agarose for imaging the periderm at 5dpf.

(D-E) Selected frames (D) and quantification (E) from live-imaging of laser-induced cellular peridermal injury of *Tg(krt4:GFP-CAAX)* zebrafish larvae either untreated or pre-treated with $Gd^{3+}$ (2.5µM, 16h). Yellow asterisk- cell selected for laser injury; blue arrowhead- junctional closure underneath extruded apoptotic cell.

(F) Stills of live imaging of laser-induced apoptosis in mosaic cultures of WT and P1-KO monolayers (extension of Fig 1-G,H). Yellow asterisk- cell selected for laser injury; magenta arrowheads- myosin ring formation at dead-neighbour cell interface, as visualised by MRLC-GFP; blue arrowhead- junctional closure underneath extruded apoptotic cell.

(G)

Untreated controls were treated with respective drugs vehicles. Scale bars: 10µm. XY panels are maximum projection views of all z-stacks. All data are means ± SEM. ***$p<0.001$, calculated from n≥3 independent experiments analysed with unpaired Student's *t* test (B), or two-way ANOVA (E).

**Figure S3.**

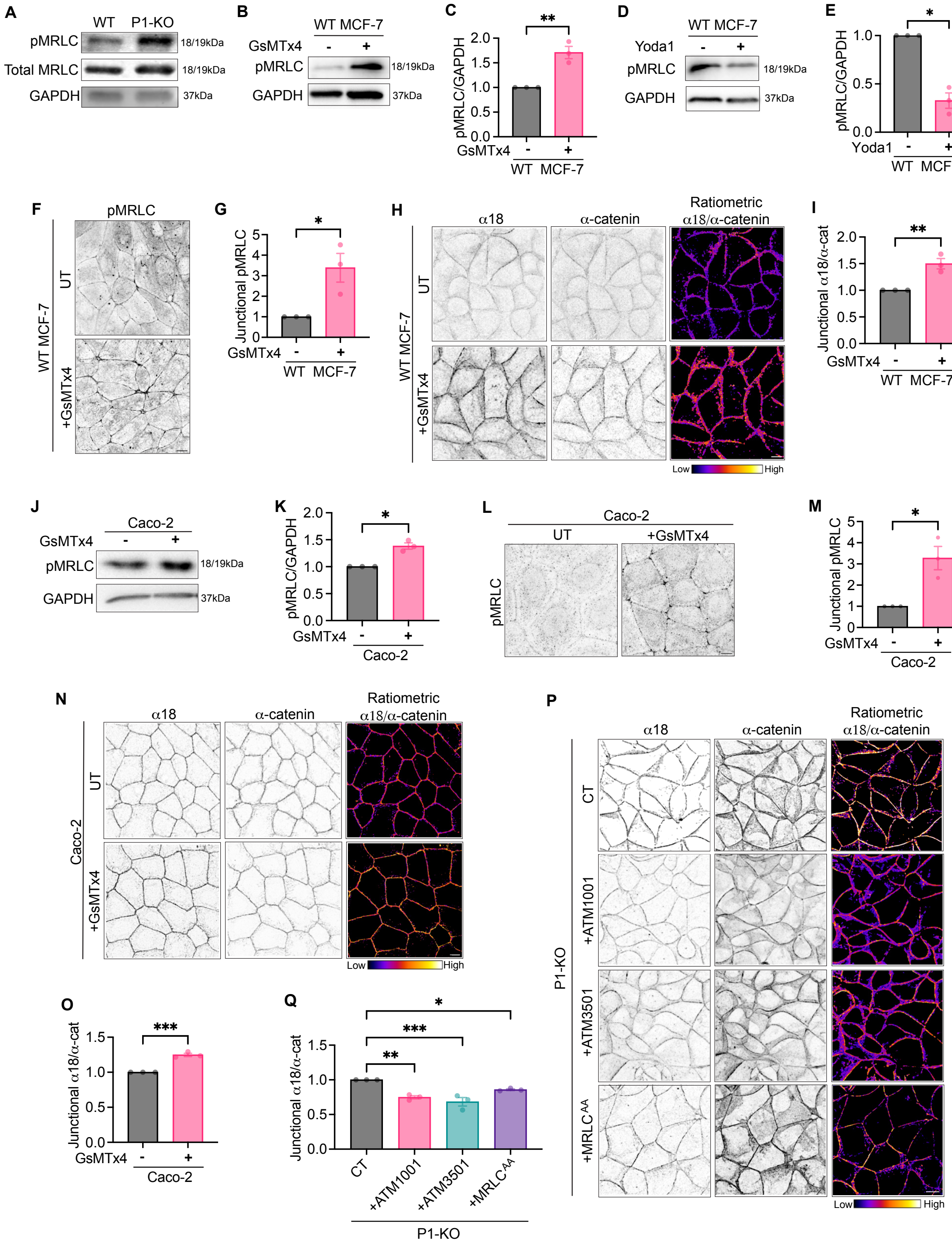


### Figure S3.

(A) Representative immunoblot of activated form of MyosinII (pMRLC) and total regulatory domain of Myosin II (MRLC) in WT and P1-KO cells.

(B-C) Representative immunoblot (B) and quantification (C) of total pMRLC levels detected in untreated and GsMTx4 (2.5μM, 15min) treated MCF7$^{WT}$ cells.

(D-E) Representative immunoblot (D) and quantification (E) of total pMRLC levels detected in untreated and Yoda1 (25μM, 15min) treated MCF7$^{WT}$ cells.

(F-G) Representative immunofluorescent images (F) and quantification (G) of pMRLC localised at junctions of untreated and GsMTx4 (2.5μM, 15min) treated MCF7$^{WT}$ monolayers.

(H-I) Representative immunofluorescent images (H) of junctional localisation of tension-sensitive α-catenin epitope (α18) and total α-catenin levels in untreated and GsMTx4 (2.5μM, 15min) treated $MCF7^{WT}$ monolayers. Ratiometric images (H) and quantification (I) represent junctional intensity of α18 normalised to total junctional α-catenin.
(J-K) Representative immunoblot (J) and quantification (K) of total pMRLC levels detected in untreated and GsMTx4 (2.5μM, 15min) treated Caco-2 cells.
(L-M) Representative immunofluorescent images (L) and quantification (M) of pMRLC localised at junctions of untreated and GsMTx4 (2.5μM, 15min) treated Caco-2 monolayers.
(N-O) Representative immunofluorescent images (N) of junctional localisation of tension-sensitive α-catenin epitope (a18) and total a-catenin levels in untreated and GsMTx4 (2.5 μM, 15min) treated Caco-2 monolayers. Ratiometric images (N) and quantification (O) represent junctional intensity of a18 normalised to total junctional α-catenin.
(P-Q) Representative immunofluorescent images (P) of junctional localisation of tension-sensitive a-catenin epitope (α18) and total α-catenin levels in monolayers of P1-KO control, treated with tropomyosin inhibitors (ATM1001, ATM3501) (2.5μM of each, 8h), and expressing phosphodeficient mutant $MRLC^{AA}$. Ratiometric images (P) and quantification (Q) represent junctional intensity of α18 normalised to total junctional α-catenin. Untreated controls were treated with respective drug vehicles. Scale bars: 10μm. XY panels are maximum projection views of all z-stacks. All data are means ± SEM. *p<0.05, **p<0.01, ***p<0.001, ****p<0.0001 calculated from n=3 independent experiments analysed with one-sample *t* test (C,E,G,I,K,M,O), or one-way ANOVA (Q).

Figure S4.

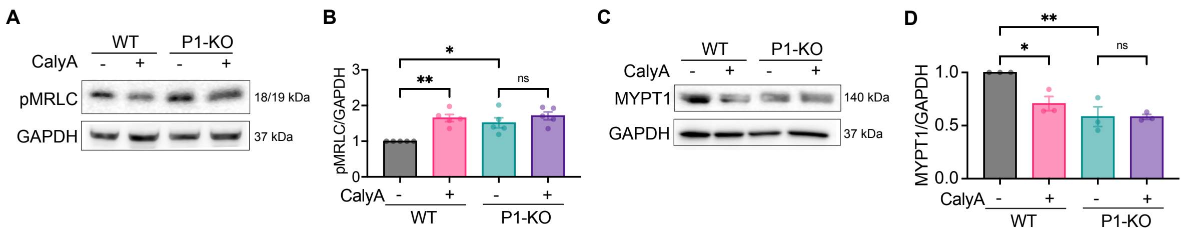


**Figure S4.**

(A-B) Representative immunoblot (A) and quantification (B) of pMRLC levels in untreated and calyculin A (CalyA, 200nM, 15min) treated WT and P1-KO cells.

(C-D) Representative immunoblot (C) and quantification (D) of MYPT1 levels in untreated and calyculin A (CalyA, 200nM, 15min) treated WT and P1-KO cells.

Untreated controls were treated with respective drug vehicles. All data are means ± SEM. *$p<0.05$, **$p<0.01$, calculated from n≥3 independent experiments analysed with or two-way ANOVA (B,D).

**Supplementary Videos**

**Video S1.** Intracellular calcium dynamics of untreated WT and P1-KO monolayers, and P1-KO monolayer treated with tropomyosin inhibitor ATM3501 (2.5μM, 8h), upon laser-injury induced apoptotic extrusion, as visualised with Calbryte-520AM dye (related to Figure S1F). Injured cell (marked with white asterisk), upon apoptosis causes spread of calcium wave across the WT monolayer, which is limited in untreated and ATM3501-treated P1-KO monolayer .

**Video S2.** Apical extrusion eliminates apoptotic $MCF7^{WT}$ cells, which is seen compromised in P1-KO monolayers (related to Figure 1B). Visualisation of contractile networks (MRLC-GFP) at interface of dying cell (yellow asterisk) and its surrounding neighbours.

**Video S3.** Apical extrusion eliminates (left) apoptotic P1-KO cell surrounded by WT cells, but is compromised when (right) P1-KO cells surround WT apoptotic cell (related to Figure S2F). Visualisation of contractile networks (MRLC-GFP) at interface of dying cell (red asterisk) and its surrounding neighbours.

**Video S4.** Apical extrusion eliminates apoptotic cells from 5-dpf zebrafish periderm (related to Figure 1E, S2D). Visualisation of lamellipodia (GFP-CAAX) in neighbours as apoptotic cell (red asterisks) is extruded from control (left) 5dpf zebrafish periderm after laser injury, but is compromised upon GsMTx4 (centre) or $Gd^{3+}$ (right) overnight pre-treatment.

**Video S5.** Apical extrusion compromised in apoptotic P1-KO cell upon laser injury, but rescued when P1-KO cells express phosphodeficient myosin mutant, $MRLC^{AA}$ (related to Figure 2N). Visualisation of contractile networks (MRLC-GFP) at interface of dying cell (red asterisk) and its surrounding neighbours.
